# Generation of stable microtubule superstructures by binding of peptide-fused tetrameric proteins to inside and outside

**DOI:** 10.1101/2022.01.27.476107

**Authors:** Hiroshi Inaba, Yurina Sueki, Muneyoshi Ichikawa, Arif Md. Rashedul Kabir, Takashi Iwasaki, Hideki Shigematsu, Akira Kakugo, Kazuki Sada, Kazunori Matsuura

## Abstract

Microtubules (MTs) play important roles in biological functions by forming superstructures, such as doublets, triplets, and branched structures, *in vivo.* Formation of these superstructures by exogenous molecules *in vitro* will be useful not only for understanding the functions of MTs but also as components of MT-based nanomaterials. Here, we developed a tetrameric fluorescent protein Azami-Green (AG) fused with a His-tag and Tau-derived peptide (TP), TP–AG, which can bind to the inside or outside of MTs depending on the polymerization conditions. The binding of TP–AG to the inside of MTs induced the formation, stabilized, and increased the rigidity of the MTs. The binding of TP–AG to the outside of MTs induced various types of MT superstructures, including doublets, multiplets, and branched structures, by recruiting tubulins to MTs. The formation of motile MT aster structures by TP–AG was also observed. The generation of MT superstructures by these exogenous proteins provides guidelines for the design of MT-based nanomaterials.

## Introduction

Microtubules (MTs) are tubular cytoskeletons composed of tubulin molecules, which are used as components of motile nanomaterials, such as active matters^1–3^ and molecular robots^4^ by complexation with motor proteins, such as kinesin. Although the MTs produced *in vitro* usually have singlet structures, various MT structures are formed *in vivo*, including structures with different lengths and protofilament numbers, and multiplets (doublets or more)^5^, and branched structures^6,7^. MT superstructures are formed as needed to perform various functions *in vivo.* For instance, flagella and cilia possess complex MT structures, including doublet and branched MTs, and these MT superstructures are thought to contribute to the mechanical strength and motility of flagella and cilia^8–10^. MT-organizing centers (MTOCs), known as the centrosomes in animal cells, contain MT nucleators to control the nucleation, stabilization, and orientation of MTs for the formation of complex MT assemblies, such as spindles and asters^11,12^. One of the ways that the structures of MTs are controlled *in vivo* is by the binding of MT-associated proteins (MAPs)^13^. Although MAPs that bind to the outer surface of MTs are well known, various MT inner proteins (MIPs) that bind to the inner surface of MTs have recently been reported^9,14–18^. The formation of doublet MTs is thought to occur by (1) the binding of MIPs to the inside of the complete A-MT for induction of a stabilized formation of an incomplete B-MT on the outer junction and (2) the binding of other proteins, such as PACRG and FAP20, to the inner junction to close the B-MT^19^. The branching of MTs is mainly known to be induced by centrosomes and the g-tubulin ring complex (g-TuRC)^20,21^. Branching structures arise by the nucleation of new MTs from the sides of existing MTs via g-TuRC and other proteins^22–28^. Although complex MT superstructures, such as the doublets and branches observed in nature, are supposed to have interesting functions distinct from singlet MTs, it is difficult to artificially construct the MT superstructures *in vitro.* Recently, the formation of doublet MTs from singlet MTs *in vitro* has been reported by deletion of the C-terminal tails of MTs to allow the access of new tubulin^29^. However, this method does not allow the branching of doublet MTs into two singlet MTs as observed in human sperm flagella^27^. Furthermore, there have been no reported examples of the generation of complicated MT structures, such as doublets and branches, by exogenous molecules. If such MT superstructures can be constructed by exogenous molecules *in vitro,* they would be useful for understanding their properties/functions *in vivo* and as new types of MT building blocks for nanotechnological applications including double-track railways^8^. For this purpose, the formation of stable MTs as a nucleus and the recruitment of new tubulin to the formed MTs are thought to be important^29^.

As a binding motif for the internal surface of MTs, we have previously developed a Tau-derived peptide (TP; CGGGKKHVPGGGSVQIVYKPVDL)^30^. TP was designed using a repeat domain of Tau that is involved in binding to the inner surface of MTs through interaction with a taxol (paclitaxcel)-binding pocket on β-tubulin^31^. The binding of TP to the interior of MTs can be achieved by the preincubation of TP with tubulin followed by the subsequent polymerization of the TP-tubulin complex. Thus, the conjugation of exogenous molecules to TP can result in their encapsulation inside MTs^14^. TP has been used to encapsulate various nano-sized materials, such as green fluorescent protein (GFP)^32^, magnetic cobalt–platinum nanoparticles^33^, and gold nanoparticles^30^ in MTs. The TP motif will be useful not only to encapsulate exogenous molecules in MTs, but also to recruit tubulins to pre-existing MTs by displaying TP on the outer surface.

Here, using the tetrameric protein Azami-Green (AG), which shows fluorescence by tetramerization^34^, we developed modified AG proteins that can bind both to the inside and outside of MTs using a simple strategy. The tetrameric structure of AG is useful to display multiple peptides on the surface for the efficient binding to MTs (Fig. 1a). By adding a His-tag to AG, AG was successfully recruited to the outer surface of MTs. TP was further fused to the C-terminus of AG (TP–AG), to display four TPs on a single tetrameric protein (Fig. 1a), allowing binding to the inner surface of tubulins. Treatment with TP–AG induced MT formation and stabilization and increased the rigidity of the MTs, enabling the formation of a variety of MT superstructures, including doublets, multiplets, branched structures, extremely long MTs, and aster structures. Because various MT structures could be generated by one type of exogenous protein, we proposed a simple design principle for forming diverse MT superstructures *in vitro.*

**Fig. 1.**
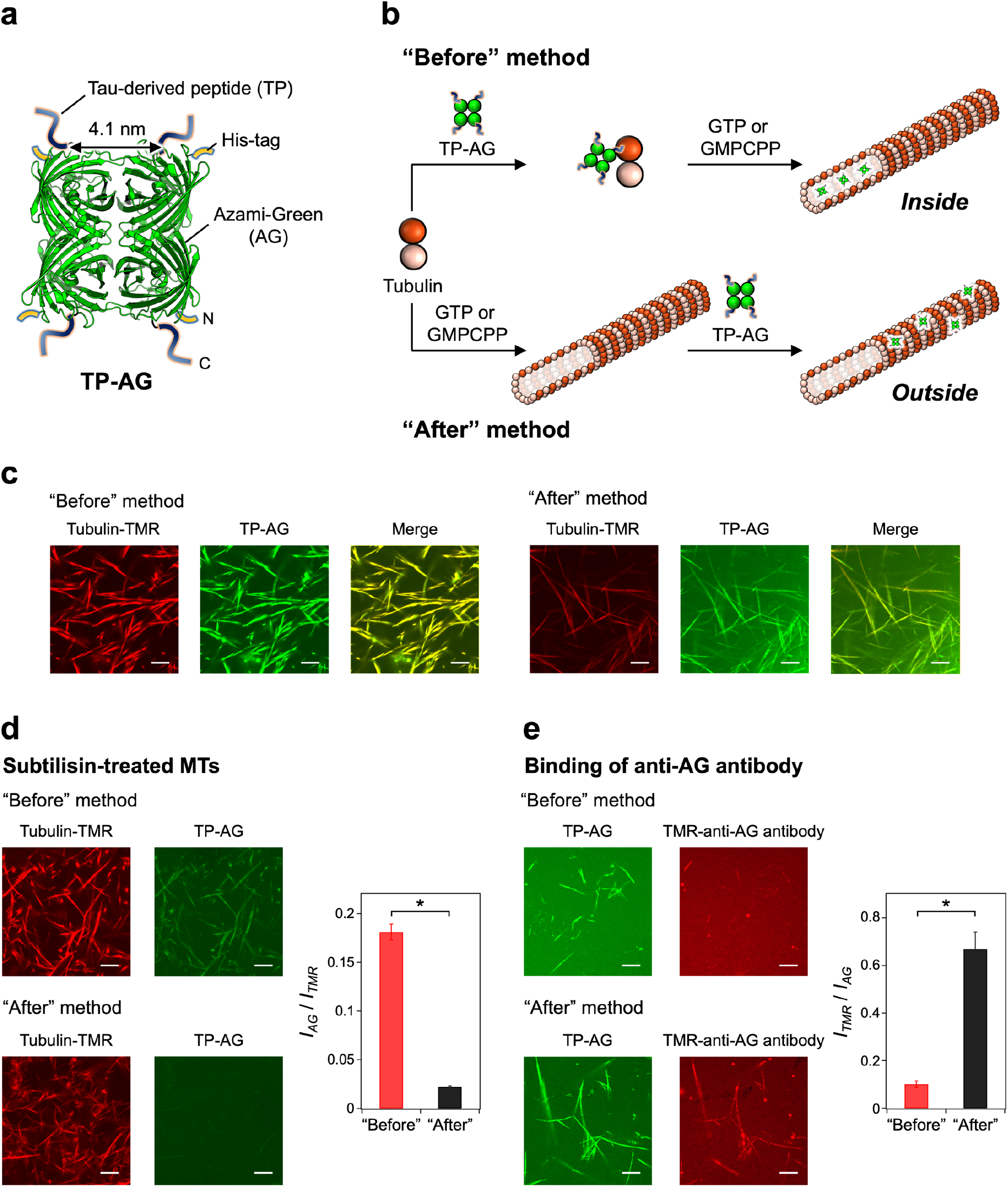
Binding of TP–AG to microtubules (MTs). **(a)** Schematic diagram of Tau-derived peptide (TP)-fused Azami-Green (AG) (TP–AG). **(b)** Preparation of TP–AG-incorporated MTs by the “Before” and “After” methods and **(c)** the confocal laser scanning microscopy (CLSM) images. Preparation concentrations: [tubulin] = 3.2 μM; [tubulin-TMR] = 0.8 μM; [TP–AG] = 8 μM; [GMPCPP] = 0.2 mM. Scale bars, 10 μm. **(d)** CLSM images of TP–AG-incorporated TMR-labeled GMPCPP MTs using subtilisin-treated tubulin prepared by the “Before” and “After” methods (left). Scale bars, 10 μm. The *I*_AG_/*I*_TMR_ ratio for each type of MT determined from the CLSM images (right). Error bars represent the standard error of the mean (*N* = 30). \**P* < 0.0001, t-test. Preparation concentrations: [tubulin] = 19.2 μM; [tubulin-TMR] = 4.8 μM; [TP–AG] = 8 μM; [subtilisin] = 1 μM; [GMPCPP] = 0.2 mM. **(e)** CLSM images of TP–AG-incorporated GMPCPP MTs prepared by the “Before” and “After” methods, and after further treatment with a TMR-labeled anti-AG antibody (left). Scale bars, 10 μm. The *I*_TMR_/*I*_AG_ ratios for each type of TP–AG-incorporated MT treated with TMR-labeled anti-AG antibody, determined from the CLSM images (right). Error bars represent the standard error of the mean (*N* = 30). \**P* < 0.0001, t-test. Preparation concentrations: [tubulin] = 4 μM; [TP–AG] = 8 μM; [TMR-labeled anti-AG antibody] = 1.2 μM; [GMPCPP] = 0.2 mM.

## Results and discussion

### Binding analysis of TP–AG to MTs

Some MT-binding proteins have higher affinities for MTs with attached His-tags because of the electrostatic interactions between the His-tags and the C-terminal tail of tubulin^35,36^. We hypothesized that adding several His-tags to non-MT-related proteins may give the proteins the ability to interact with MTs. Thus, we attached a His-tag to the tetrameric AG protein so that there would be four His-tags per tetrameric AG (Supplementary Table 1 and Supplementary Fig. 1). Although AG without a His-tag does not have the ability to interact with MTs^37^, the binding of His-tagged AG (referred to as AG hereafter for simplicity) to MTs prepared with guanosine-5’-[(α,β)-methyleno]triphosphate (GMPCPP) (GMPCPP MTs) was observed by confocal laser scanning microscopy (CLSM) through the colocalization of AG and tetramethylrhodamine (TMR)-labeled MTs (Supplementary Fig. 2a, top) and by the co-sedimentation of AG and MTs (Supplementary Fig. 2b). Treatment with subtilisin, which digests the *C*-terminal tails of tubulin located on the outer surface of MTs^29,38^, suppressed the binding of AG to the MTs (Supplementary Fig. 2a, bottom). This result indicated that AG was bound to the C-terminal tails on the outer surface of the MTs, presumably thorough the interaction between the His-tags of tetrameric AG and the C-terminal tails of tubulin. In addition, the attachment of AG to MTs prepared with GTP (GTP MTs) and numerous bundling structures of AG-treated GMPCPP MTs were observed in negative-stain electron microscopy (EM), showing the linking of MTs by the binding of AG to the outer surface of MTs (Supplementary Fig. 2c).

AG with His-tags was observed to bind to the outer surface of the MTs, and therefore we attached the TP sequences to the AG to form TP–AG (Fig. 1a and Supplementary Table 1). The binding of TP–AG to MTs was evaluated by two methods: the “Before” and “After” methods (Fig. 1b). In the “Before” method, tubulin was first incubated with TP–AG, and then MTs were formed by the addition of GTP or GMPCPP. In the “After” method, TP–AG was incubated with preassembled MTs. In the present study, GMPCPP was mainly used for analysis of the binding of TP–AG and the effects on the MT structures, and GTP was mainly used for analysis of the effects of TP–AG on the formation and stability of the MTs. The binding of TP–AG to GMPCPP MTs was observed in both the “Before” and “After” methods by CLSM (Fig. 1c). We analyzed the binding ratio of TP–AG to GMPCPP MTs by a co-sedimentation assay and found that almost all TP–AG was bound to MTs at high concentrations of tubulin, with *K_d_* = 11 μM in the “Before” method (Supplementary Fig. 3). Unlike for AG, high binding of TP–AG to MTs was observed in the “Before” method when subtilisin-treated tubulin was used, whereas the binding was low in the “After” method (Fig. 1d). In addition, the binding of a TMR-labeled anti-AG antibody to TP–AG-incorporated MTs was low in the “Before” method, whereas more binding was observed when the construct was prepared by the “After” method (Fig. 1e). These results showed that most of the TP–AG was bound to the inner surface of MTs in the “Before” method and to the outer surface of MTs in the “After” method (Fig. 1b), similar to that observed for TP-conjugated GFP^32^. We also constructed TP–mAG with TP linked to a monomeric version of AG (mAG)^34^ (Supplementary Table 1 and Supplementary Fig. 1). The binding of TP–mAG to MTs was much weaker than for tetrameric TP–AG in both the “Before” and “After” methods (Supplementary Fig. 4). Thus, the tetrameric structure of TP–AG is important for the binding to MTs and the binding of TP–AG to the inside or outside of the MTs can be controlled by changing the polymerization method.

### Effects of TP–AG on the polymerization and stability of MTs

Since GTP MTs are typically unstable *in vitro* due to the hydrolysis of GTP to GDP, the effects of TP–AG on the polymerization and stability of GTP MTs were analyzed. A considerable increase in the turbidity of the tubulin solution with TP–AG suggested that addition of TP–AG caused a dramatic enhancement in MT formation (Fig. 2a). TP–AG was much more effective in facilitating MT formation compared with the MT-stabilizing anticancer drug, taxol. MT formation was not facilitated by AG or TP–mAG, indicating that both the tetramerization of TP–AG and the binding of TP–AG to the inside of the MTs were necessary for the increased induction of MT formation. The formation efficiency of GTP MTs prepared by the “Before” method was evaluated by SDS-PAGE of the separated free tubulin and MTs (Fig. 2b). Increased formation of MTs was observed by the treatment of TP–AG, and the MT structures were partially retained even after treatment at 4°C for 90 min (depolymerization conditions). GTP MTs with TP–AG prepared by the “Before” method were observed to be stable with longer lengths compared with taxol-stabilized MTs, and only aggregates were observed after treatment with AG or without any additives (Fig. 2c). To examine the effects of TP–AG on the physical stability of the MTs, EM was performed (Fig. 2d). With the TP sequence attached, the MT bundling effect of AG was reduced compared with AG without TP (Supplementary Fig. 2c). In the negative-stain EM images, the MTs with TP–AG had a straighter appearance compared with normal MTs, which appeared to have a winding shape (Fig. 2d, left), indicating that TP–AG physically strengthened the MT structures. This result was confirmed by analyzing the ratio of the contour and end-to-end lengths of the MTs (Fig. 2d, right). These results indicated that the binding of TP–AG to the inside of the MTs formed rigid and stable MTs, and considerably promoted MT formation. The MTs treated with TP–AG prepared by the “After” method also showed straighter structures compared with the winding structures of normal MTs (Fig. 2d). It is possible that the binding of TP–AG to the inner and outer surfaces had similar strengthening effects on the MTs or that the partial binding of TP–AG to the outer surface in the “Before” method was enough to strengthen the MTs.

**Fig. 2.**
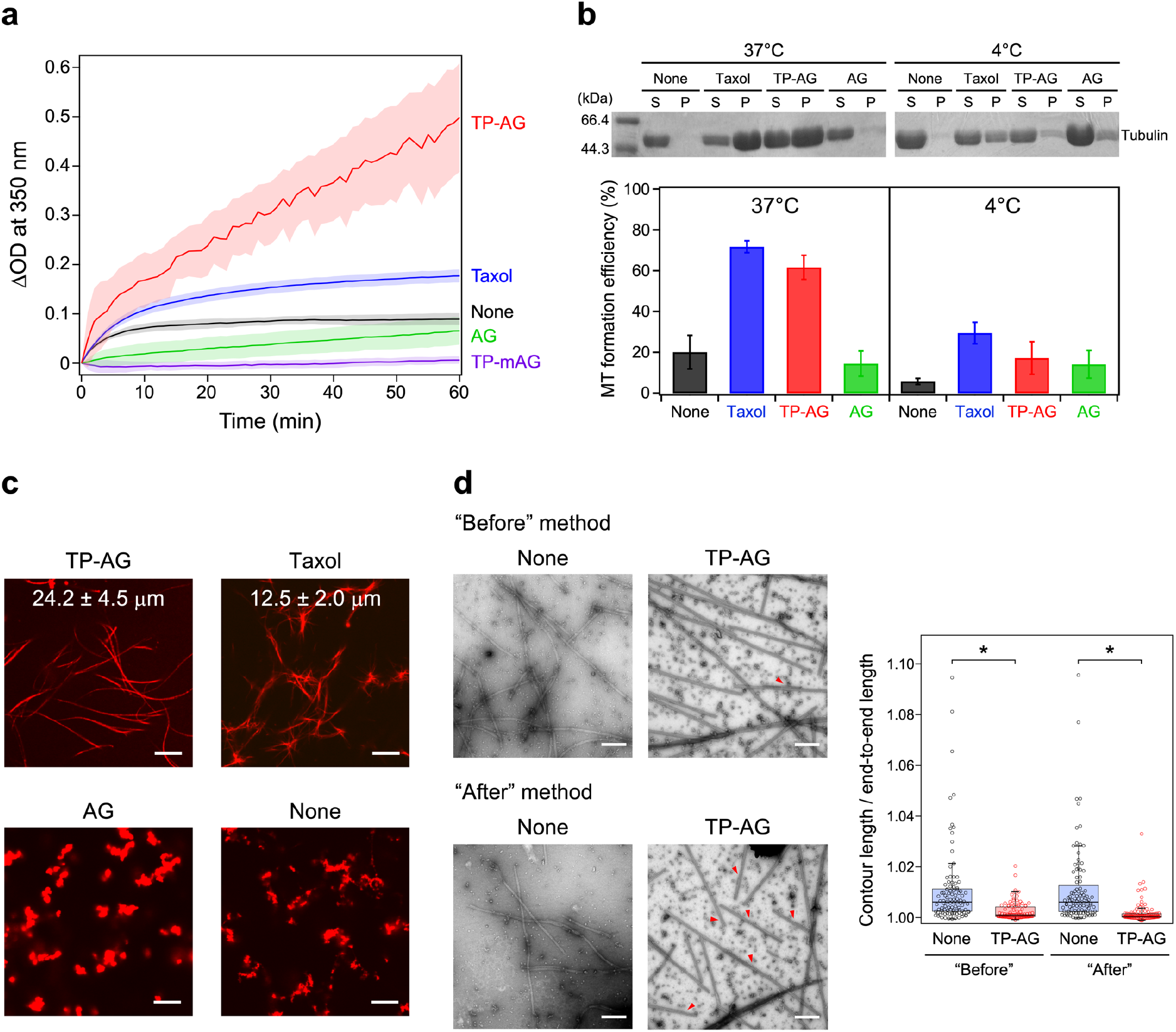
Facilitation of the polymerization and stabilization of MTs by TP–AG. **(a)** Changes in turbidity over time caused by tubulin polymerization. The optical density at 350 nm was measured for 4 μM tubulin and 1 mM GTP in the presence of 10 μM TP–AG (red), AG (green), TP–mAG (purple), taxol (blue), or without any additives (black) at 37°C. The thick lines represent the average values and the colored areas show the standard errors of the mean of three independent experiments. **(b)** Effect of TP–AG on the formation of GTP MTs prepared by the “Before” method. Preparation concentrations: [tubulin] = 30 μM; [TP–AG] = 34 μM; [AG] = 34 μM; [taxol] = 34 μM; [GTP] = 1 mM. Samples were prepared by polymerization at 37°C for 30 min (37°C) and by polymerization at 37°C for 30 min and subsequent depolymerization at 4°C for 90 min (4°C). SDS-PAGE results of the supernatants (S) and the pellets (P) obtained by centrifugation of the solutions of the mixtures (top). The MT formation efficiency as calculated by the ratio of S and P (bottom). Error bars represent the standard error of the mean of three independent experiments. **(c)** CLSM images of GTP MTs prepared by the “Before” method with TP–AG, taxol, AG, and without any additives. Scale bars, 10 μm. The lengths of the MTs (average ± standard deviation, *N* = 30) are shown. Preparation concentrations: [tubulin] = 3.2 μM; [tubulin–TMR] = 0.8 μM; [TP–AG] = 5 μM; [AG] = 5 μM; [taxol] = 5 μM; [GTP] = 1 mM. **(d)** Effect of TP–AG on the straightness of GMPCPP MTs prepared by the “Before” and “After” methods evaluated by negative-stain electron microscopy (EM) (left). Red arrowheads, doublet MTs. Scale bars, 500 nm. Box plot of the ratios of the contour and end-to-end lengths of MTs obtained from the negative-stain EM images (*N* = 100) (right). *\*P* < 0.0001, t-test. Preparation concentrations: [tubulin] = 10 μM; [TP–AG] = 20 μM; [GMPCPP] = 0.2 mM.

### MT superstructures induced by TP–AG

In the negative-stain EM observations of TP–AG-treated GMPCPP MTs, there were doublet-like MTs and branched MTs present, not just singlet MTs, especially in the “After” method (Fig. 2d and Supplementary Fig. 5a). With further incubation at 37°C for 30 min in the “After” method, the populations of doublet MTs (44.0%) and branched MTs (5.6%) were increased in the negative-stain EM images (Fig. 3a and 3b) compared with the MTs without further incubation (Fig. 3b). There were also regions where multiplets were formed (Fig. 3a, orange arrowheads) and the formation of multiplet and branched structures was taking place at the same time (Fig. 3a, green arrowheads). The doublet MTs, branched MTs, and multiplets were also observed by cryo-EM (Fig. 3c and Supplementary Fig. 5b). The structures of the doublet MTs induced by TP–AG were similar to the doublet MTs isolated from cilia (Supplementary Fig. 5c) and doublet MTs constructed *in vitro*^29^, which consisted of complete singlet MTs (A-MTs) and incomplete MTs surrounding AMTs (B-MTs). Because most MTs (97.1%) were singlet MTs in the absence of TP–AG and the population of the doublet MTs with TP–AG in the “Before” method (9.0%) was less than in the “After” method (Fig. 3b), it was thought that these doublet-like MTs were formed via TP–AG bound to the outside of the preassembled MTs. From the EM images, the branching was caused by the B-MT detaching from the A-MT of doublet MTs. The method of the branching was different from that caused by the nucleation factor SSNA1 in which the tubulin lattice of one singlet MT is separated into two singlet MTs^22^. The amount of branched MTs was increased by further incubation at 37°C for 30 min, supporting the concept that the branched structures were grown from doublet MTs under the polymerization conditions (Fig. 3b). Because the lengths of the B-MTs of the doublet MTs were similar with (0.47 ± 0.25 μm) and without (0.40 ± 0.18 μm) incubation (Fig. 3d), when the B-MTs were partially grown, the MTs might split into two separate singlets rather than keep growing as a doublet. Because the doublet MTs formed by subtilisin treatment did not form branched structures^29^, it is possible that the remaining C-terminal tails of tubulin under the conditions used in the present study pushed the formed B-MT off and thereby branched structures grew from the doublet MTs. The branches had a mean angle of 4.9 ± 2.5° (Fig. 3e) and this small angle was similar to the branched MTs that are split from doublet MTs in cilia^27^ rather than the branches induced by SSNA1, which have wider angles (47 ± 15°)^22^. Only singlet MTs were observed when the MTs were treated with AG only (Supplementary Fig. 2c), indicating the TP moiety was required for the formation of doublet and branched MTs. In a previous report, the digestion of the C-terminal tails of MTs by subtilisin made free tubulin molecules accessible to induce the formation of doublet MTs^29^. In our system, it is plausible that TP–AG bound to the outer surface of the MTs covered the C-terminal tails of the MTs, and then the exposed TP moieties recruited free tubulin to induce the formation of doublet MTs (Fig. 3f). The tetrameric structure of TP–AG with four His-tags and four TPs is possibly important for the efficient binding to the outer surface of MTs and the recruitment of tubulin. *In vivo,* one study has proposed that different types of MIPs may work together to stabilize MTs and shield the C-terminal tail of tubulin to generate doublet MTs^29^. The formation of doublet and branched MTs with TP–AG in the presence of the C-terminal tail is a model of the formation of doublet and branched MTs *in vivo.* Notably, we were able to achieve this result using a single type of protein.

**Fig. 3.**
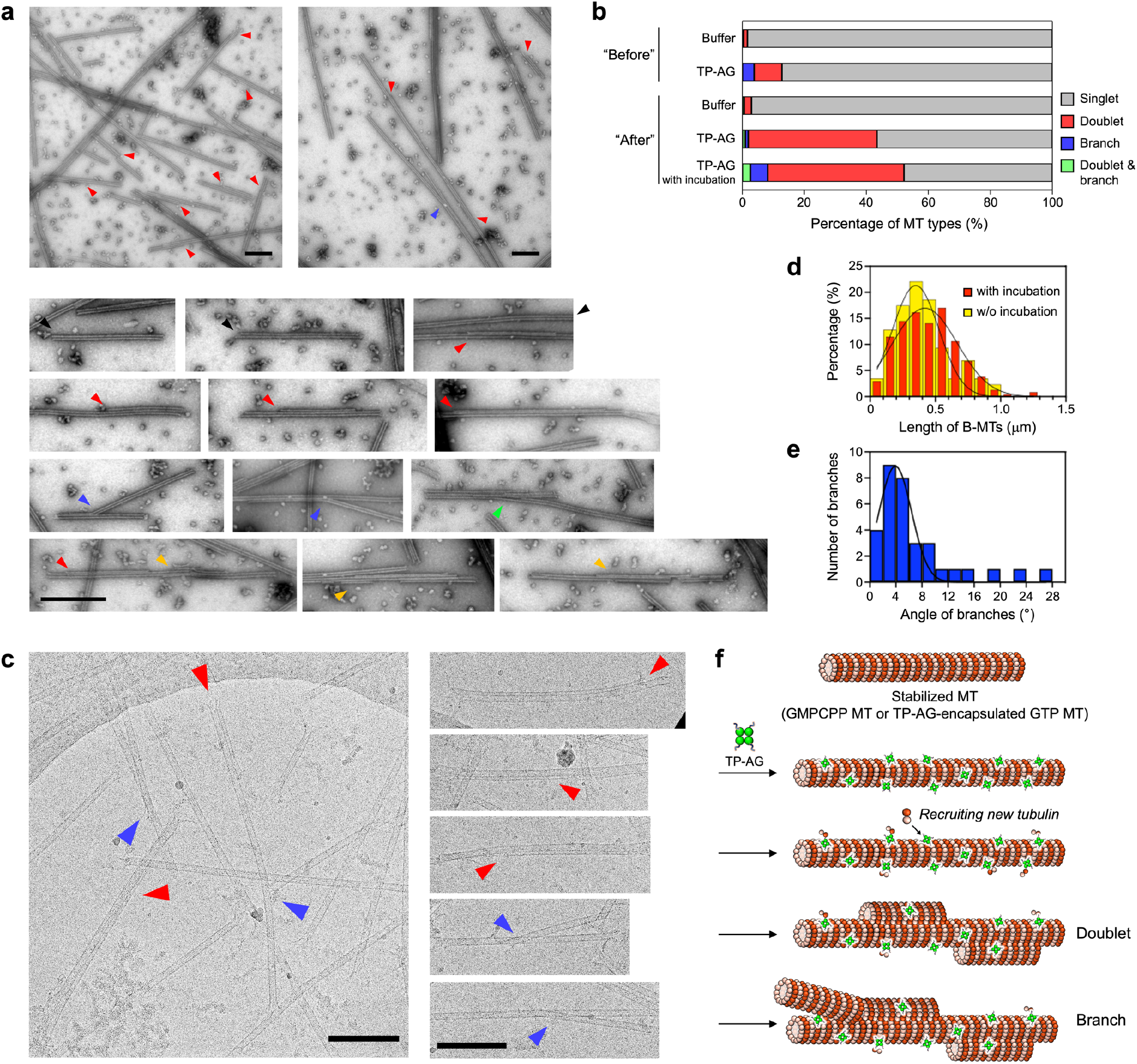
Various MT structures induced by TP–AG. **(a)** Negative-stain EM images of TP–AG-incorporated GMPCPP MTs prepared by the “After” method with additional incubation at 37°C for 30 min. General view (upper row) and selected EM images (lower rows) of MTs are shown. When TP–AG was added to preassembled MTs, various MT structures: singlet MTs (black arrowheads); doublet MTs (red arrowheads); branched MTs (blue arrowheads); and multiplet MTs (orange arrowheads) were observed. The green arrowhead indicates the location where branching and the formation of multiplets are happening in the same area. Scale bars, 500 nm. **(b)** Quantification of MT types in GMPCPP MTs prepared by the “Before” method treated with buffer (98.4% singlet, 1.2% doublet, 0.4% branched, and 0% doublet and branched MTs, *N* = 248) and TP–AG (87.2% singlet, 9.0% doublet, 3.8% branched, and 0% doublet and branched MTs, *N* = 290), and by the “After” method treated with buffer (97.1% singlet, 2.3% doublet, 0% branched, and 0.6% doublet and branched MTs, *N* = 171), TP–AG (56.6% singlet, 41.5% doublet, 0.9% branched, and 0.9% doublet and branched MTs, *N* = 212), and TP–AG with further incubation at 37°C for 30 min (47.9% singlet, 44.0% doublet, 5.6% branched, and 2.6% doublet and branched MTs, *N* = 539) from the negative-stain EM images. Multiplet MTs are counted as “Doublet” here. Preparation concentrations: [tubulin] = 10 μM; [TP–AG] = 20 μM; [GMPCPP] = 0.2 mM. **(c)** Representative cryo-EM images of TP–AG-incorporated GMPCPP MTs prepared by the “After” method with additional incubation at 37°C for 30 min. A general view is shown in the left panel and each type of MT is shown in the right column. Red arrowheads, doublet MTs. Blue arrowheads, branched MTs. Similar MT structures to those observed with negative-stain EM were also observed using cryo-EM. Scale bars, 250 nm. **(d)** Histogram of the lengths of the B-MTs of doublet MTs with (red) and without (yellow) incubation at 37°C for 30 min. The curves are Gaussian fits to the histogram (with incubation: mean 0.47 μm and s.d. 0.25 μm, *N* = 235; without incubation: mean 0.40 μm and s.d. 0.18 μm, *N* = 86). **(e)** Histogram of the distribution of the angles of the branched MTs. The curve is a Gaussian fit to the histogram (mean 4.9° and s.d. 2.5°, *N* = 33). **(f)** Model for the formation of doublet and branched MTs with TP–AG.

The effect of TP–AG on GTP MTs was also assessed by negative-stain EM. When 10 μM tubulin was used, only aggregates were observed in the absence of TP–AG (Fig. 4a). Increasing the amount of TP–AG in the “Before” method increased the amount of MTs formed (Fig. 4b–f). When the amount of TP–AG was low, the majority of the MTs formed were singlet MTs, whereas complex structures, such as doublet MTs, multiplet MTs, and branched MTs, were observed by increasing the equivalents of TP–AG in the presence of GTP (Fig. 4g and 4h). The formation of doublet MTs may require a certain level of stabilization of the MTs because it has been reported that doublet MTs were formed only when stable GMPCPP MTs were used and not when unstable GTP MTs were used^29^. In our system, first TP–AG binds to the inner surface of MTs to stabilize singlet MTs, then additional TP–AG binds to the outer surface to induce the formation of doublet MTs. When the ratio of tubulin to TP–AG was increased to 1:4, extremely long and winding MTs were observed (Fig. 4h). This result was consistent with the CLSM images in which TP–AG encapsulated MTs were longer than taxol-stabilized MTs (Fig. 2c). These extremely long MTs are thought to be formed by a combination of the formation of a doublet (or multiplet) MT, splitting of the doublet, and the weak bundling activity of TP–AG. This complex MT structure is thought to lead to the high stability of the TP–AG-treated MTs.

**Fig. 4.**
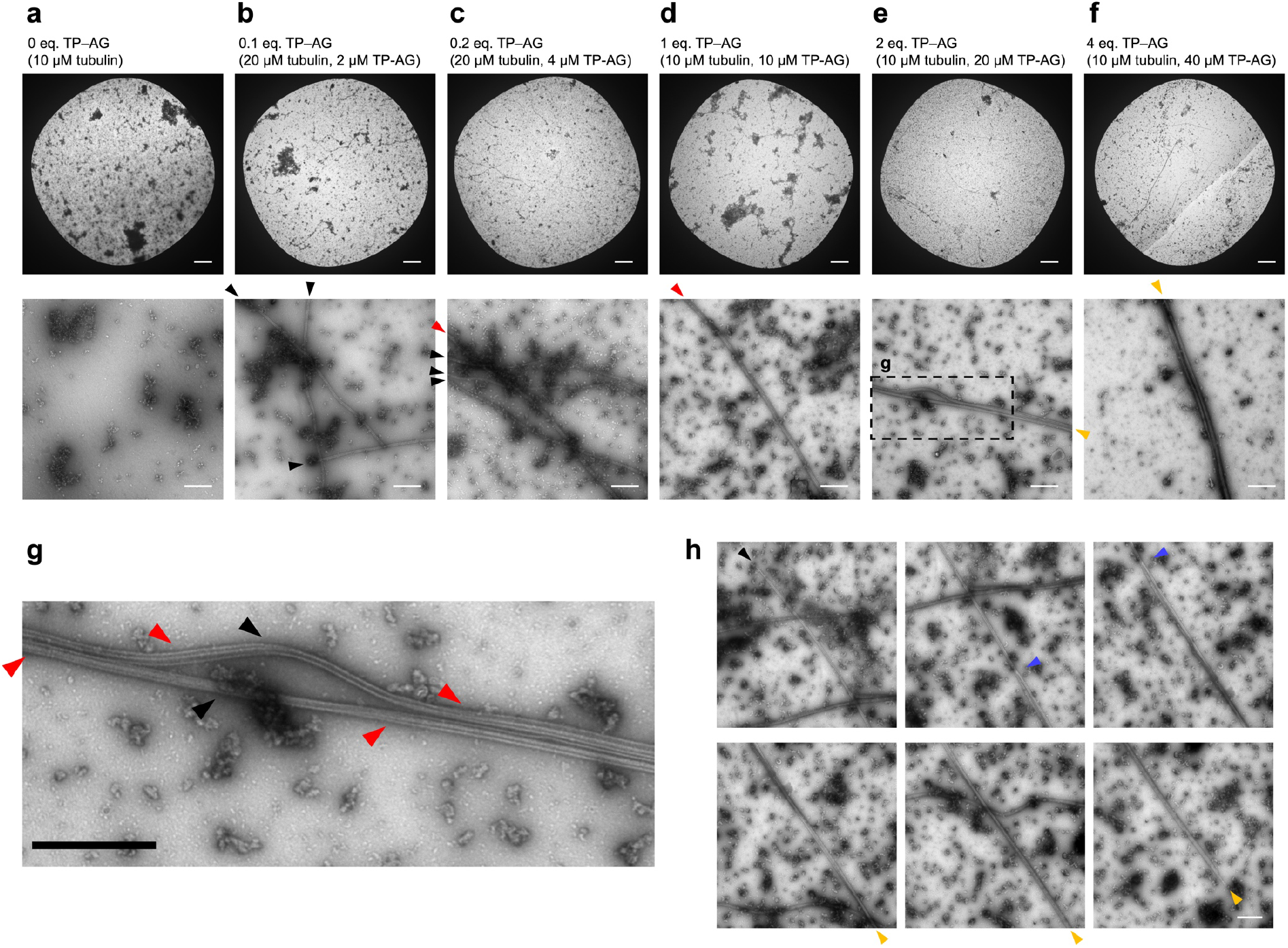
Extremely long MTs induced by TP–AG. **(a)-(f)** Representative negative-stain EM images of GTP MTs treated with TP–AG by the “Before” method with increasing amounts of TP–AG as indicated above. The upper row shows lower magnification with the carbon hole and the bottom row shows higher magnification EM images. Without TP–AG, 10 μM tubulin alone did not yield MTs (**a**). With a lower concentration of TP–AG, the majority of the MTs were singlet MTs (black arrowheads). When higher concentrations of TP–AG were added, more complicated MT structures were formed, including doublet MTs (red arrowheads) and multiplet MTs (orange arrowheads). The inset (dashed line) indicates the location of (**g**). Scale bars, 10 μm for top and 500 nm for bottom images. **(g)** Detail of multiplet MT formation. Even within one MT, there were both singlet areas (black arrowheads) and doublet areas (red arrowheads). Scale bar, 500 nm. **(h)** Example of a branched MT. The MT was prepared with 2 equivalents TP–AG as in (**e**). EM image of one continuous MT structure with overlapping areas. A singlet MT (black arrowhead) was split into two singlets at the locations indicated by the blue arrowheads. These two singlet MTs were intertwined together (orange arrowheads). Scale bar, 500 nm.

### Motile properties of TP–AG-incorporated MTs

The *in vitro* motility of MTs driven by ATP on a kinesin-coated substrate is the transformation of chemical energy into mechanical motion, which is fundamental property of MTs for constructing dynamic systems such as cargo delivery, active matters, and molecular robots^1–4^. To investigate the effects of TP–AG to the motility of MTs, the gliding properties of MTs treated with TP–AG were evaluated (Fig. 5a). The velocity of TP–AG-incorporated GMPCPP MTs prepared by the “Before” method was increased (1.3-fold) compared with normal MTs. The persistence length (*L*_p_) of the MTs, an indicator of the rigidity, was also increased (3.3-fold) with TP–AG, which corresponded with the negative-stain EM images (Fig. 2d). Because rigid GMPCPP MTs move faster than flexible GTP MTs^39^, it was assumed that the increased rigidity of the TP–AG-incorporated MTs was responsible for the enhanced velocity. The formation of aster-like structures composed of the GTP MTs treated with 0.2 equivalents of TP–AG to tubulin was observed in the “Before” method (Fig. 5b and 5c). At the center of the asters, a strong accumulation of TP–AG was observed and the centers showed no motility. Accumulated MTs with less TP–AG at the periphery of the asters showed motility that was released and translocated away from the aster. This radially organized motility from the asters was similar to that observed with the asters from MTOCs reconstituted from *Xenopus* sperm^40^. Construction of MT aster structures has been reported using MT nucleating structures, such as DNA origami^41^ and beads coated with antibodies to Aurora kinase A, which plays a key role in MT nucleation^43,43^. It is possible that the MTs with accumulated TP–AG in the present study serve as MT nucleating centers to trap MTs with less TP–AG, followed by the release of the trapped MTs from the center. To our knowledge, this is a first example that a single kind of exogenous protein works as a nucleating center. When an increased amount of TP–AG was added, aster structures were not formed and bundle-like structures with no motility were formed instead (Supplementary Fig. 6). Thus, we were able to form different MT superstructures by balancing the content of TP–AG-incorporated and non-TP–AG incorporated MTs.

**Fig. 5.**
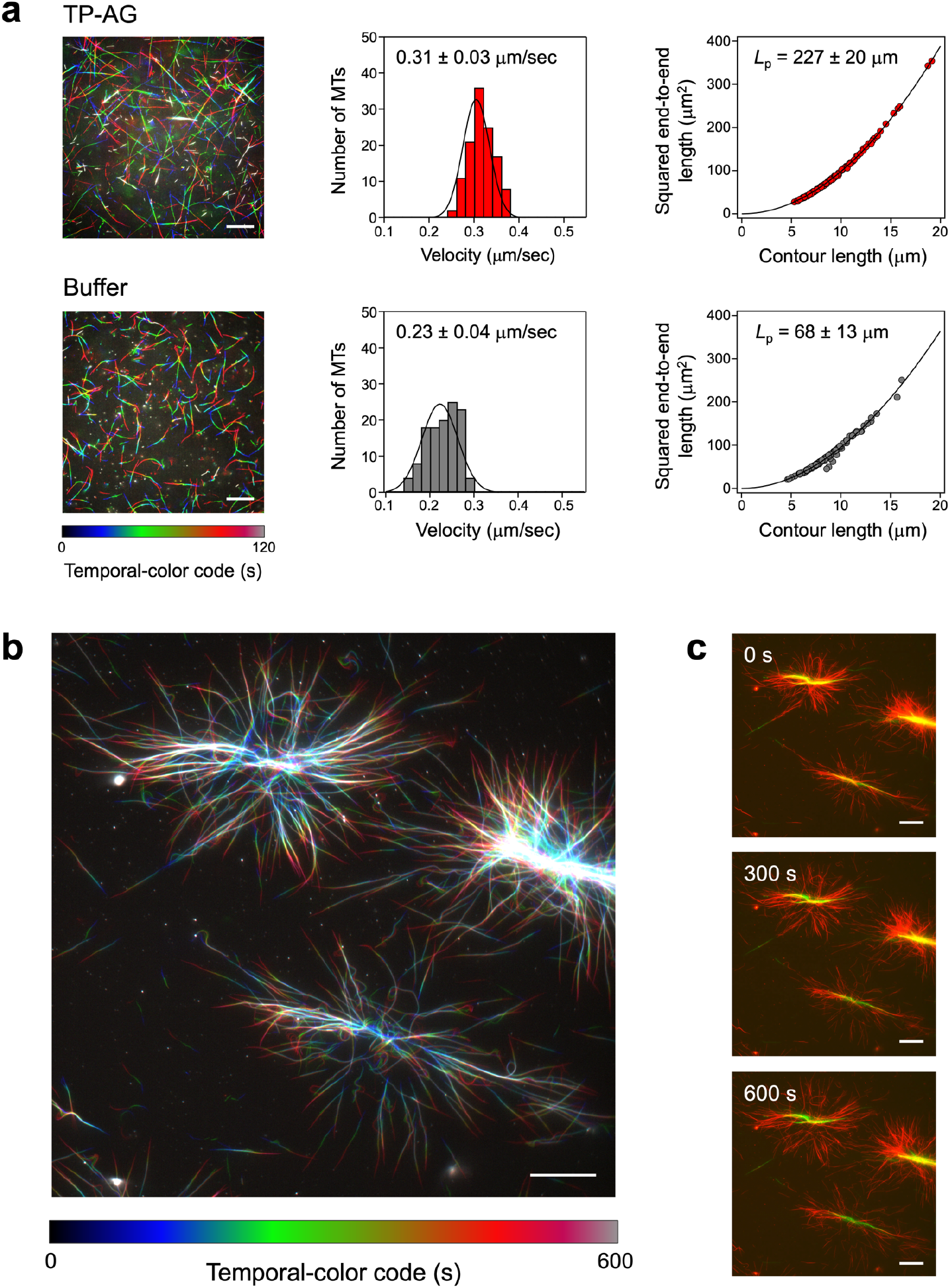
Motile properties of TP–AG-incorporated MTs. **(a)** Effect of TP–AG on the velocity and persistence length (*L*_p_) of GMPCPP MTs prepared by the “Before” method on a kinesin-coated substrate. Fluorescence microscopy image of moving MTs using a temporal-color code (left). Histogram of the velocity of the MTs (center, *N* = 120). The curves are Gaussian fits to the histogram. *L*_p_ of MTs estimated using the curves that are derived from equation (3) in Methods (right, *N* = 120). Data presented as average ± standard deviation. The difference in velocities and *L*_p_ of MTs treated with TP–AG and buffer was found to be statistically significant (*P* < 0.0001, t-test). Preparation concentrations: [tubulin] = 6.4 μM; [tubulin–TMR] = 1.6 μM; [TP–AG] = 0.8 μM; [GMPCPP] = 0.2 mM. See also Supplementary Movie 1. **(b)** Fluorescence microscopy image of moving aster structures of TP–AG-incorporated GTP MTs prepared by the “Before” method on a kinesin-coated substrate using a temporal-color code. To visualize the movement of MTs in (**a**) and (**b**), the time series images were color coded and superimposed on one another. The color scales are shown at the bottom of the image. **(c)** Time-lapse images showing the motility of the aster structures of TP–AG-incorporated GTP MTs. Preparation concentrations: [tubulin] = 4 μM; [tubulin–TMR] = 1 μM; [TP–AG] = 1 μM; [GTP] = 1 mM. See also Supplementary Movie 2. Scale bars, 20 μm.

## Conclusions

Using a simple design based on fusing a His-tag and TP to exogenous tetrameric AG, we succeeded in stabilizing MTs and inducing the formation of various MT superstructures, including doublets, multiplets, branched MTs, and extremely long MTs, by binding TP–AG to both the inside and outside of MTs. In addition, the formation of MTOC-like motile aster structures was achieved. These results provide design guidelines for the recruitment of exogenous proteins to MTs and the construction of MT superstructures. Highly stable, rigid, and long MT superstructures are useful for MT-based nanotechnological applications, such as active matters, molecular robots, double-track railways, and reconstructed shafts^44^. These results also provide important insights into the formation principles of doublets and branched structures *in vivo*, such as in flagella and cilia.

## Acknowledgments

This work was supported by JSPS KAKENHI Grant Numbers JP19K15699 to H. I., JP20K15733 and JP21KK0125 to M. I., JP20H05972 and JP21K04846 to A.M.R.K., JP18H05423 and 21H04434 to A.K. from the Japan Society for the Promotion of Science (JSPS), ACT-X (JPMJAX2012 for H. I.), FOREST Program (JPMJFR2034 for H. I.), and PRESTO (JPMJPR20E1 for M. I.) from the Japan Science and Technology Agency (JST), and Future AI and Robot Technology Research and Development Project from the New Energy and Industrial Technology Development Organization (NEDO), Japan (for A.K.). We thank technical stuffs Hitomi Ichikawa and Atsunobu Suzuki from Nara Institute of Science and Technology for the help of EM observation. We thank Victoria Muir, PhD, from Edanz (https://jp.edanz.com/ac) for editing a draft of this manuscript.

## Author contributions

H.I., Y.S., and M.I. contributed equally to this work. H.I. and K.M. conceived the original idea and H. I. designed the AG proteins. T.I. constructed the plasmids of the AG proteins, expressed the AG proteins, and helped to purify the AG proteins. Y.S. purified the AG proteins, evaluated the binding to MTs and effects on the polymerization and stability of MTs, and performed the motility assay. M.I. measured negative-stain EM. M.I. and H.S. measured cryo-EM. A.M.R.K., A.K., and K.S. purified tubulin. A.M.R.K. purified kinesin, prepared TP–AG-incorporated MTs and aster MT structures, and observed their motility in the motility assay. H.I., M.I., and K.M. wrote the entire paper and all authors revised the entire paper. H.I., M.I., and T.I. wrote the experimental details.

## Methods

### Equipment and materials

UV-vis spectra were obtained using a Jasco V-630. Ultracentrifugation was performed using an Optima MAX-TL ultracentrifuge (Beckman Coulter) using TLA 120.2 rotor. Confocal laser scanning microscopy (CLSM) measurement was carried out using a FluoView FV10i (Olympus). In the motility assay, samples were illuminated with a LED light source and visualized by using an epi-fluorescence microscope (Eclipse Ti2-E; Nikon) using an oil-coupled Lambda S 60x objective (NA 1.4) (Nikon). For the observation of aster structures, samples were illuminated with a 100 W mercury lamp and visualized by using an epi-fluorescence microscope (Eclipse Ti; Nikon) using an oil-coupled Plan Apo 60x objective (NA 1.4) (Nikon). Tubulin was purified from porcine brain by a reported procedure^45^. Tetremethylrhodamine (TMR)-labeled tubulin (tubulin–TMR) was prepared following a standard protocol^46^. Recombinant kinesin-1 consisting of the first 573 amino acid residues of human kinesin-1 was prepared according to a reported procedure^47^. In the SDS-PAGE analysis, the proteins were mixed with 2×Laemmli buffer, heated at 95°C for 5 min, then loaded onto SDS-PAGE (15% acrylamide). The reagents used were purchased from Watanabe Chemical Ind., Ltd., Tokyo Chemical Industry Co., Dojindo Laboratories Co., Ltd., Wako Pure Chemical Industries, and Sigma-Aldrich. All the chemicals were used without further purification. For negative-stain EM, holey carbon girds (U1013, EMJapan Co.,Ltd.) were pre-hydrophilized using an E-1010 ion sputter (Hitachi). Negatively stained grids were observed in an H-7100 electron microscope (Hitachi) equipped with AMT XR41 digital CCD camera (Advanced Microscopy Techniques). For cryo-EM, holey carbon grids (Quantifoil R2/1 grid: Cu 300mesh, M2951C-1-300) were hydrophilized by a JEC-3000FC Auto Fine Coater (JEOL) with a target of Aluminum. Vitrification of the grids were performed using a Vitrobot Mark IV (Thermo Fisher Scientific), and vitrified grids were observed in a Glacios cryo transmission electron microscope (cryo-TEM) (Thermo Fisher Scientific) equipped with Falcon III direct electron detector (Thermo Fisher Scientific) at SPring-8. Concentrations of TP–AG and AG are described as monomer concentrations.

### Construction of plasmids

The TP–AG, AG, and TP–mAG expression vectors were constructed based on the pET-29b(+) expression vector (Merck, Darmstadt, Germany). The TP–AG, AG, and TP–mAG genes were synthesized to optimize codon usage for *Escherichia coli*. The open circular pET-29b(+) vector was synthesized by inverse polymerase chain reaction (PCR). PCR was performed in the TaKaRa PCR Thermal Cycler Dice Touch (TaKaRa, Shiga, Japan) using the Tks Gflex DNA Polymerase (TaKaRa). The synthetic genes coding TP–AG, AG, and TP–mAG were ligated into the open circular pET-29b(+) using the In-Fusion HD Cloning Kit (TaKaRa) to generate the recombinant protein expression vectors. The plasmids were sequenced by Eurofins Genomics (Tokyo, Japan).

### Expression and purification of proteins

The pET-29b(+) vectors coding TP–AG, AG, and TP–mAG were transformed into *E. coli* BL21(DE3) strain by a heat-shock procedure. Bacterial cells were spread on Luria-Bertani– Ampicillin (100 μg/mL) (LBA) agar and grown overnight at 37°C. A single transformant colony was grown in LBA medium at 37°C overnight. The culture was diluted 100-fold by addition to fresh LBA medium and grown at 37°C until an absorbance of 0.5 was noted at 600 nm (mid-logarithmic phase), and then the culture was incubated with 0.1 mM isopropyl β-D-1-thiogalactopyranoside at 20°C. After 18 h of incubation, cells were harvested by centrifugation at 8000 rpm for 10 min. The cell pellets were suspended in Ni-affinity binding buffer (50 mM Tris-HCl pH 7.4, 150 mM NaCl, 20 mM imidazole) on ice. The cells were lysed by sonication. After centrifugation at 13000 rpm for 10 min, the supernatant was loaded onto 1 mL Ni-affinity column (GE healthcare). After washing with the same buffer and the storage buffer (50 mM Tris-HCl pH 8.0, 300 mM NaCl), the protein was eluted from the column by using the Ni-affinity elution buffer (50 mM Tris-HCl pH 8.0, 300 mM NaCl, 250 mM imidazole). The eluted sample was dialyzed against the storage buffer (Spectra/por7, cutoff Mw: 8 kDa, Spectrum Laboratories, Inc.) at 4°C. The purity of proteins was evaluated by SDS-PAGE (Supplementary Fig. 1). The concentration of proteins was determined using reported molar extinction coefficients^34^.

### Construction of TP–AG-incorporated GMPCPP MTs

In the typical “Before” method, TP–AG (10 μM) was mixed with a solution containing tubulin (4 μM) and tubulin–TMR (1 μM) in BRB80 (80 mM PIPES pH 6.9, 1.0 mM MgCl_2_, 1.0 mM EGTA) and incubated at 25°C for 30 min in the dark. Then, 2 μL of GMPCPP premix (1 mM GMPCPP, 20 mM MgCl_2_ in BRB80) was added to the mixture (8 μL) and incubated at 37°C for 30 min in the dark (Final concentrations: [Tubulin] = 3.2 μM, [Tubulin–TMR] = 0.8 μM, [TP–AG] = 8 μM, [GMPCPP] = 0.2 mM). As a control, AG or TP–mAG (8 μM) was used instead of TP–AG.

In the typical “After” method, GMPCPP premix (2 μL) was mixed with a solution (6 μL) containing tubulin (4 μM) and tubulin–TMR (1 μM) in BRB80. The mixture was incubated at 37°C for 30 min in the dark. Then TP–AG (2 μL) was added to the mixture and kept at 25°C for 30 min in the dark (Final concentrations: [Tubulin] = 3.2 μM, [Tubulin–TMR] = 0.8 μM, [TP–AG] = 8 μM, [GMPCPP] = 0.2 mM). As a control, AG or TP–mAG (8 μM) was used instead of TP–AG.

### Removal of C-terminal tails of tubulin with subtilisin

C-terminal tails of tubulin were removed by the treatment of subtilisin according to the reported procedure with modification^29^. Mixture of tubulin (64 μM) and tubulin-TMR (16 μM) were treated with subtilisin in a subtilisin:tubulin ratio of 1:100 (w/w) at 4°C for 30 min. Then the subtilisin activity was inhibited by incubation with 12.5 mM PMSF (phenylmethylsulphonyl fluoride) at 4°C for 10 min. After centrifugation at 21000 rpm at 4°C for 20 min, the supernatants were collected and used as subtilisin-treated tubulin. The removal of C-terminal tails of tubulin were confirmed by SDS-PAGE (Supplementary Fig. 2a).

### Binding analysis of TP–AG to GMPCPP MTs using subtilisin-treated tubulin

TP–AG-incorporated GMPCPP MTs were prepared by the “Before” or “After” method as above using subtilisin-treated tubulin or untreated tubulin (Final concentrations: [Tubulin] = 19.2 μM, [Tubulin–TMR] = 4.8 μM, [TP–AG] = 8 μM, [GMPCPP] = 0.2 mM). As a control, AG (8 μM) was used instead of TP–AG. The mixture was used for CLSM imaging. Fluorescence intensity per MT was calculated from the fluorescence images by subtracting the background intensity using ImageJ software (NIH). The background-subtracted AG fluorescence intensity per TMR fluorescence intensity for each MT (*N* = 30) were calculated (Fig. 1d).

### Preparation of TMR-labeled anti-AG antibody

Anti-AG polyclonal antibody (1.3 mg/mL, 190 μL) in PBS (137 mM NaCl, 8.1 mM Na_2_HPO_4_, 2.68 mM KCl, 1.47 mM KH_2_PO_4_) was mixed with 5-carboxytetramethylrhodamine, succinimidyl ester (10 mM, 10 μL) in DMSO. The mixture was incubated at 37°C for 15 min in the dark. The TMR-labeled anti-AG antibody was obtained by washing with PBS seven times by the ultrafiltration (cutoff Mw: 10 kDa). The concentrations of TMR and the anti-AG antibody of the TMR-labeled antibody were determined by UV-vis spectrometry. The modification ratio of TMR per the anti-AG antibody was determined as 2.2.

### Binding analysis of the anti-AG antibody to TP–AG-incorporated GMPCPP MTs

TP–AG-incorporated GMPCPP MTs were prepared by the “Before” or “After” method as above. Then the TMR-labeled anti-AG antibody (6 μM, 2 μL) prepared above was added to the TP–AG-incorporated MT solution (8 μL) and incubated at 25°C for 1 h in the dark (Final concentrations: [Tubulin] = 4 μM, [TP–AG] = 8 μM, [TMR-labeled anti-AG antibody] = 1.2 μM, [GMPCPP] = 0.2 mM). The mixture was used for CLSM imaging. Fluorescence intensity per MT was calculated from the fluorescence images by subtracting the background intensity using ImageJ. The background-subtracted TMR fluorescence intensity per AG fluorescence intensity for each MT (*N* = 30) were calculated (Fig. 1e).

### Binding analysis of TP–AG to GMPCPP MTs by co-sedimentation assay

TP–AG-incorporated GMPCPP MTs were prepared by the “Before” method as above (Final concentrations: [Tubulin] = 5, 10, 20, and 30 μM, [TP–AG] = 37 μM, [GMPCPP] = 0.2 mM). As a control, AG or TP–mAG (37 μM) was used instead of TP–AG. After ultracentrifugation of the mixture at 50000 rpm at 37°C for 5 min, the supernatants and the pellets were collected and analyzed by SDS-PAGE. The ratio of TP–AG bound to MTs (*θ*) was calculated as the ratio of the density of TP–AG bands in the supernatants and the pellets. The results were treated with the following Hill equation **(1)** to give the dissociation constant (*K_d_*) and the Hill coefficient (*n*) (Supplementary Fig. 3c).

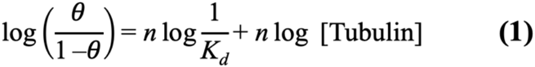

By the fitting, *K_d_* and *n* were determined as 11 μM and 2.2, respectively.

### Construction of TP–AG-incorporated GTP MTs

In the typical “Before” method, TP–AG (6.25 μM) was mixed with a solution containing tubulin (4 μM) and tubulin–TMR (1 μM) in BRB80 and incubated at 25°C for 30 min in the dark. Then, 4 μL of GTP premix (5 mM GTP, 20 mM MgCl_2_ in BRB80, 25% DMSO) was added to the mixture (16 μL) and incubated at 37°C for 30 min in the dark (Final concentrations: [Tubulin] = 3.2 μM, [Tubulin–TMR] = 0.8 μM, [TP–AG] = 5 μM, [GTP] = 1 mM). As a control, AG or taxol (5 μM) was used instead of TP–AG.

### Evaluation of the effect of TP–AG on the formation of GTP MTs

TP–AG-incorporated GTP MTs were prepared by the “Before” method as above (Final concentrations: [Tubulin] = 30 μM, [TP–AG] = 34 μM, [GTP] = 1 mM). As a control, taxol (34 μM), AG (34 μM), or the storage buffer was used instead of TP–AG. For the “37°C” samples, the samples were polymerized by incubating at 37°C for 30 min upon addition of GTP premix. For the “4°C” samples, the samples were polymerized by incubating at 37°C for 30 min upon addition of GTP premix and then further incubated at 4°C for 90 min. After ultracentrifugation at 50000 rpm at 37°C for 5 min for the “37°C” samples and at 35000 rpm at 4°C for 5 min for the “4°C” samples, the supernatants and the pellets were collected and analyzed by SDS-PAGE. The MT formation efficiency was calculated as the ratio of the density of tubulin bands in the supernatants (*S*) and the pellets (*P*) by using the following equation **(2)**.

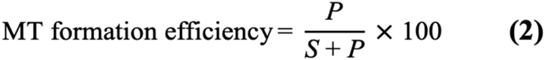

The average and the standard error of the mean of three independent measurements are shown in Fig. 2b.

### CLSM measurement

The glass bottom dishes (Matsunami, Osaka, Japan) were coated by 1 mg/mL poly-L-lysine (Mw: 30000–70000, Sigma) at room temperature for 1 h, then removed and dried. The MT samples were put on the plate and kept at room temperature for 0.5–1 h, then observed by CLSM. Tubulin–TMR was excited with 550 nm and observed through a 574 nm emission band-pass filter (Red). TP–AG, AG, and TP–mAG were excited with 493 nm and observed through a 505 nm emission band-pass filter (Green).

### Turbidity measurement

Turbidity measurements were performed with tubulin (4 μM) and GTP (1 mM) in the absence or presence of TP–AG, AG, TP–mAG, or Taxol (10 μM) in BRB80 at 37°C. The optical density at 350 nm was monitored with a UV-Vis spectrometer for 60 min at 1 min intervals. When TP–AG, AG, and TP–mAG were used, the absorbance of these proteins at 350 nm was subtracted. The average and the standard error of the mean of three independent measurements are shown in Fig. 2a.

### Negative-stain EM measurement

Samples (3.5 μL) with different polymerization methods were applied to pre-hydrophilized carbon grids and negatively stained with 1.5% uranyl acetate. The specimens were examined in an H-7100 electron microscope (Hitachi) operating at 75 kV. EM images were recorded by AMT XR41 digital CCD camera (Advanced Microscopy Techniques). MT lengths and angles of branches were measured by ImageJ using EM images taken at a nominal magnification of 30kx. To evaluate the straightness of MTs, EM images were taken at a nominal magnification of 20kx. A homemade ImageJ macro was used to obtain the coordinates of start and end points of one MT, and segmented length of the same MT at the same time. The end-to-end distances of MTs were calculated and the ratio with the segmented lengths of corresponding MTs were obtained. To analyze the proportions of the MT types, EM images were taken at a nominal magnification of 30kx. Visual inspection was performed to categorize MT types. For both measurement of straightness and analysis of proportions of MT types, MTs with at least 500-nm long were used.

### Cryo-EM measurement

For cryo-EM, polymerized GMPCPP MTs (10 μM) were incubated with TP–AG (20 μM) at 25°C for 30 min and further incubated at 37°C for at least 30 min. A total of 3.5 μL of TP–AG treated MT samples was applied to a glow discharged holey carbon grid (Quantifoil R2/1). The grids were blotted, and plunged into liquid ethane using the Vitrobot Mark IV (Thermo Fisher Scientific) at 30°C and 100% humidity with a blot force of 3 and a blot time of 5 s. The grids were observed in the Glacios cryo-TEM (Thermo Fisher Scientific) operating at 200 kV. EM images were taken with Falcon III direct electron detector (Thermo Fisher Scientific) in a linear mode with the pixel size of 4.0 Å/pixel. Total dose was ~18 electrons/Å^2^, and the defocus was set to −2.0 μm.

### Motility assay

TP–AG-incorporated GMPCPP MTs were prepared by the “Before” method as above (Final concentrations: [Tubulin] = 6.4 μM, [Tubulin–TMR] = 1.6 μM, [TP–AG] = 0.8 μM, [GMPCPP] = 0.2 mM). BRB80 was used instead of TP–AG as a control (buffer). The MTs were diluted 16-fold by BRB80. Flow cells were prepared by making a narrow channel on a 24 mm × 60 mm coverslip covered with 18 mm × 18 mm coverslip (Matsunami, Osaka, Japan) using double-sided tape as a spacer. Firstly, 0.5 mg/mL casein in BRB80 was introduced into the flow cells and incubated for 3 min. Then the solution was exchanged with Wash buffer (0.5 mg/mL casein, 4.5 mg/mL D-glucose, 50 U/mL glucose oxidase, 50 U/mL catalase, 1.0 mM DTT, 1.0 mM MgCl_2_ in BRB80) containing 800 nM kinesin and incubated for 3 min. After washing with Wash buffer, the solution was exchanged with MT solution and incubated for 3 min. After washing with Wash buffer, the solution was exchanged with Wash buffer containing 5.0 mM ATP and 1.0 mM Trolox. Then the motility of MTs was imaged every 10 sec. All the experiments were performed at room temperature. Temporal-color coded images of MT movement were generated by superimposing 13 images using the Temporal-Color Code plugin in the Fiji image processing software package based on ImageJ.

### Image analysis for motility assay

Movies of the motility assays of MTs and images obtained by the fluorescence microscopy were analyzed to determine the velocity, end-to-end length, and contour length of each MT by using ImageJ. The contour length along each MT and the end-to-end distance of the same MT were measured. Persistence length (*L*_p_) was determined by fitting the data to equation **(3)** using Excel and Solver.

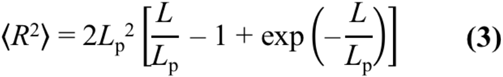

where <*R*^2^> is the mean squared end-to-end distance and *L* is the contour length^48^.

### Observation of aster structures

TP–AG-incorporated GTP MTs were prepared by the “Before” method as above (Final concentrations: [Tubulin] = 20 μM, [Tubulin–TMR] = 10 μM, [TP–AG] = 4 μM, [GTP] = 1 mM). The MTs were diluted 80-fold by a solution containing 1 mM GTP, 4 mM MgCl_2_ in BRB80, 5% DMSO. Flow cells were prepared by making a narrow channel on a 24 mm × 60 mm coverslip covered with 18 mm × 18 mm coverslip (Matsunami, Osaka, Japan) using double-sided tape as a spacer. Firstly, 0.5 mg/mL casein in BRB80 was introduced into the flow cells and incubated for 3 min. Then the solution was exchanged with Wash buffer containing 800 nM kinesin and incubated for 3 min. After washing with Wash buffer, the solution was exchanged with the diluted MT solution and incubated for 3 min. After washing with Wash buffer, the solution was exchanged with Wash buffer containing 5.0 mM ATP, 1 mM GTP, 4 mM MgCl_2_, and 5% DMSO. Then the motility of MTs was imaged every 30 sec. All the experiments were performed at room temperature. A temporal-color coded image of MT movement was generated by superimposing 21 images using the Temporal-Color Code plugin in the Fiji image processing software package based on ImageJ.

## Notes

### Competing Interest Statement

The authors have declared no competing interest.

