## Supplementary Information for "Generation of stable microtubule superstructures by binding of peptide-fused tetrameric proteins to inside and outside"

[1]Department of Chemistry and Biotechnology, Graduate School of Engineering, Tottori University, Tottori 680-8552, Japan. [2]Centre for Research on Green Sustainable Chemistry, Tottori University, Tottori 680-8552, Japan. [3]Division of Biological Science, Graduate School of Science and Technology, Nara Institute of Science and Technology, Ikoma, Nara 630-0192, Japan. [4]PRESTO, Japan Science and Technology Agency, Kawaguchi, Japan. [5]Faculty of Science, Hokkaido University, Sapporo 060-0810, Japan. [6]Department of Bioresources Science, Graduate School of Agricultural Sciences, Tottori University, Tottori 680-8553, Japan. [7]RIKEN SPring-8 Center, 1-1-1 Kouto, Sayo, Hyogo 679-5148, Japan. [8]Graduate School of Chemical Sciences and Engineering, Hokkaido University, Sapporo 060-0810, Japan.

[a]H.I., Y.S., and M.I. contributed equally to this work.

**Supplementary Table 1.** Amino acid sequences of the AG proteins used in this study.

| Protein | Sequence |
|---|---|
| AG | MHHHHHHGSMVSVIKPEMKIKLCMRGTVNGHNFVIEGEGKGNPYEGTQI<br>LDLNVTEGAPLPFAYDILTTVFQYGNRAFTKYPADIQDYFKQTFPEGYH<br>WERSMTYEDQGICTATSNISMRGDCFFYDIRFDGVNFPPNGPVMQKKTL<br>KWEPSTEKMYVRDGVLKGDVNMALLLEGGGHYRCDFKTTYKAKKDVRLP<br>DYHFVDHRIEILKHDKDYNKVKLYENAVARYSMLPSQAK |
| TP–AG | MHHHHHHGSMVSVIKPEMKIKLCMRGTVNGHNFVIEGEGKGNPYEGTQI<br>LDLNVTEGAPLPFAYDILTTVFQYGNRAFTKYPADIQDYFKQTFPEGYH<br>WERSMTYEDQGICTATSNISMRGDCFFYDIRFDGVNFPPNGPVMQKKTL<br>KWEPSTEKMYVRDGVLKGDVNMALLLEGGGHYRCDFKTTYKAKKDVRLP<br>DYHFVDHRIEILKHDKDYNKVKLYENAVARYSMLPSQAKGGGSGGGKKH<br>VPGGGSVQIVYKPVDL |
| TP–mAG | MHHHHHHGSMVSVIKPEMKIKLCMRGTVNGHNFVIEGEGKGNPYEGTQI<br>LDLNVTEGAPLPFAYDILTTVFQYGNRAFTKYPADIQDYFKQTFPEGYH<br>WERSMTYEDQGICTATSNISMRGDCFFYDIRFDGTNFPPNGPVMQKKTL<br>KWEPSTEKMYVRDGVLKGDVNMALLLEGGGHYRCDFKTTYKAKKDVRLP<br>DAHKVDHRIEILKHDKDYNKVKLYENAVARYSMLPSQAKGGGSGGGKKH<br>VPGGGSVQIVYKPVDL |

The green indicates AG or mAG and blue indicates TP. Mutated sites in mAG are shown as red.

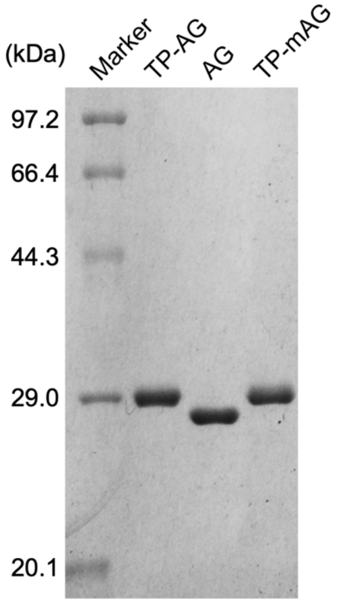


**Supplementary Figure 1.** SDS-PAGE of purified TP–AG (29.6 kDa), AG (27.2 kDa), and TP–mAG (29.5 kDa).

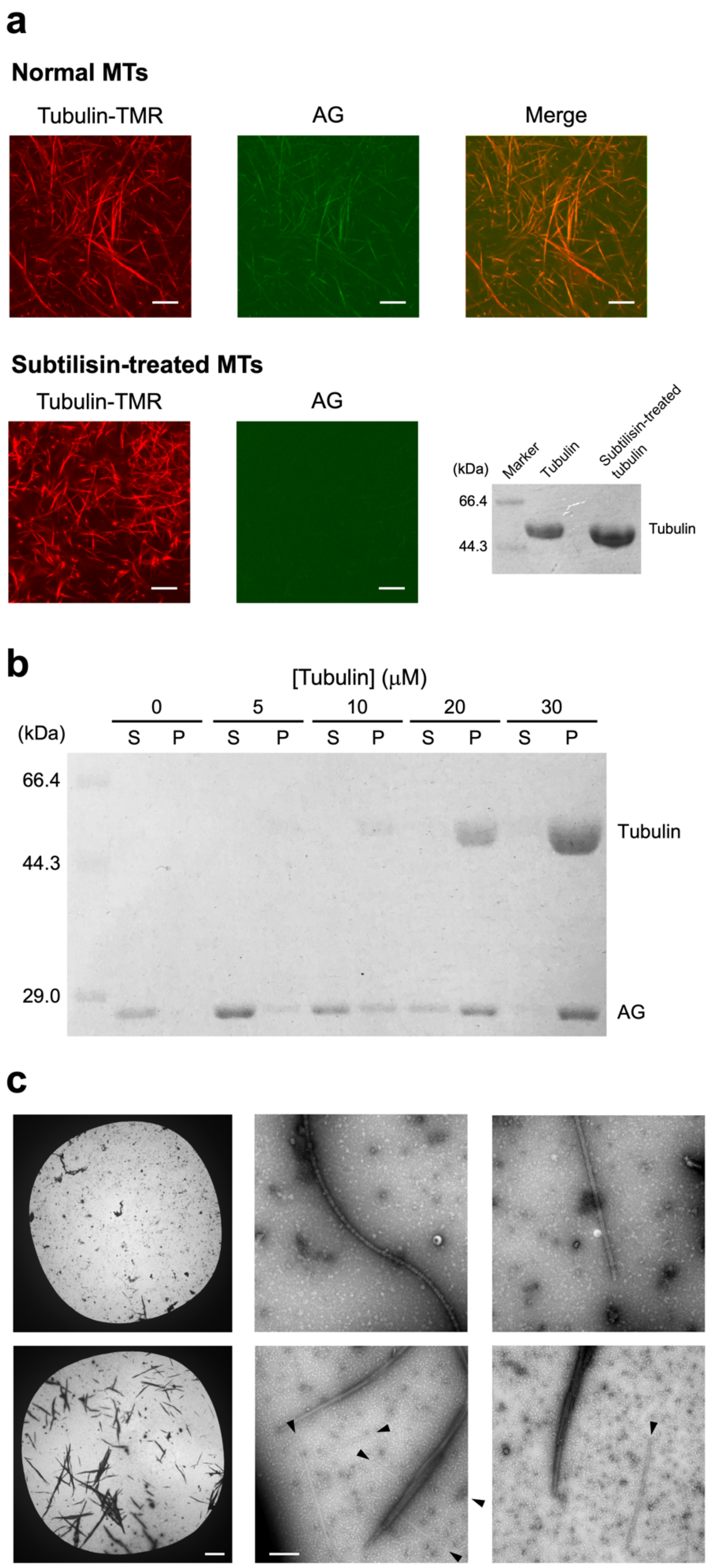


**Supplementary Figure 2.** Binding analysis of AG to MTs. **(a)** CLSM images of GMPCPP MTs incubated with AG ("After" methods). Normal MTs without subtilisin treatment or subtilisin-treated MTs were used. Preparation concentrations: [tubulin] = 19.2 μM; [tubulin-TMR] = 4.8 μM; [AG] = 8 μM; [subtilisin] = 1 μM; [GMPCPP] = 0.2 mM. SDS-PAGE results of tubulin and subtilisin-treated tubulin in a subtilisin:tubulin ratio of 1:100 (w/w). Partial removal of C-terminal tails of tubulin by the subtilisin treatment was confirmed. **(b)** Binding assay of AG to GMPCPP MTs. MTs were prepared using 37 μM AG and various concentrations of tubulins by the "Before" method. SDS-PAGE results of the supernatants (S) and the pellets (P) obtained by centrifugation of the mixture solutions is shown. **(c)** Representative negative-stain EM images of AG-treated GTP MTs (top) and GMPCPP MTs (bottom) prepared by the "Before" method. Scale bars, 10 μm for left and 500 nm for center and right. Black arrowheads, singlet MTs.

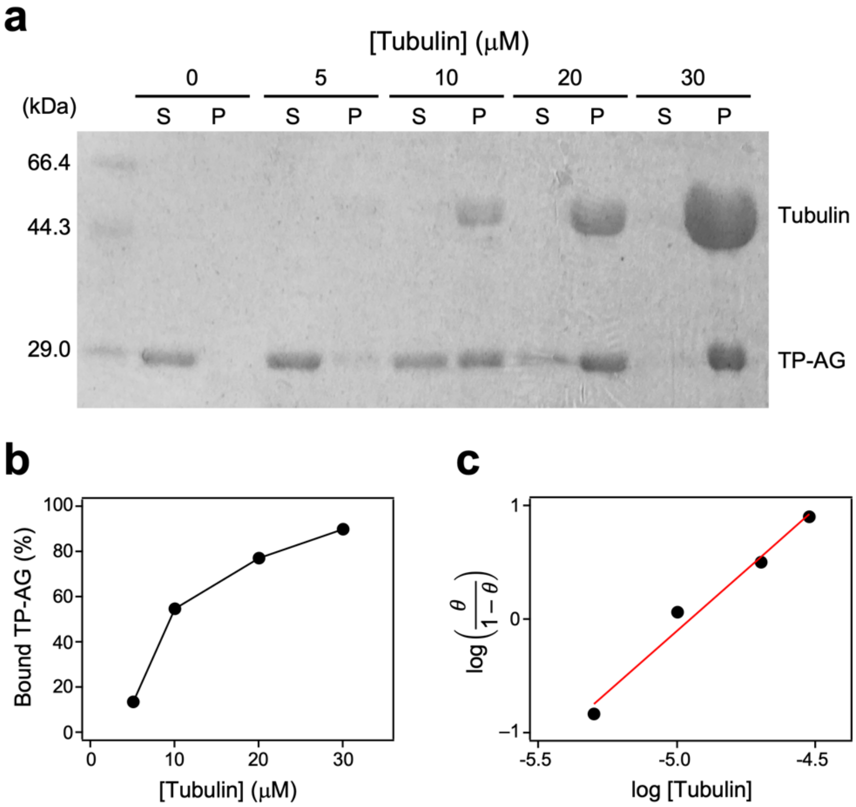


**Supplementary Figure 3.** Binding assay of TP–AG to GMPCPP MTs. MTs were prepared using 37 μM TP–AG and various concentrations of tubulins by the "Before" method. **(a)** SDS-PAGE results of the supernatants (S) and the pellets (P) obtained by centrifugation of the mixture solutions. **(b)** Percentages of TP–AG bound to MTs and **(c)** the Hill plot. $\theta$ indicates the bound ratio of TP–AG to MTs. Closed circles are experimental values and the red line is the fitted line.

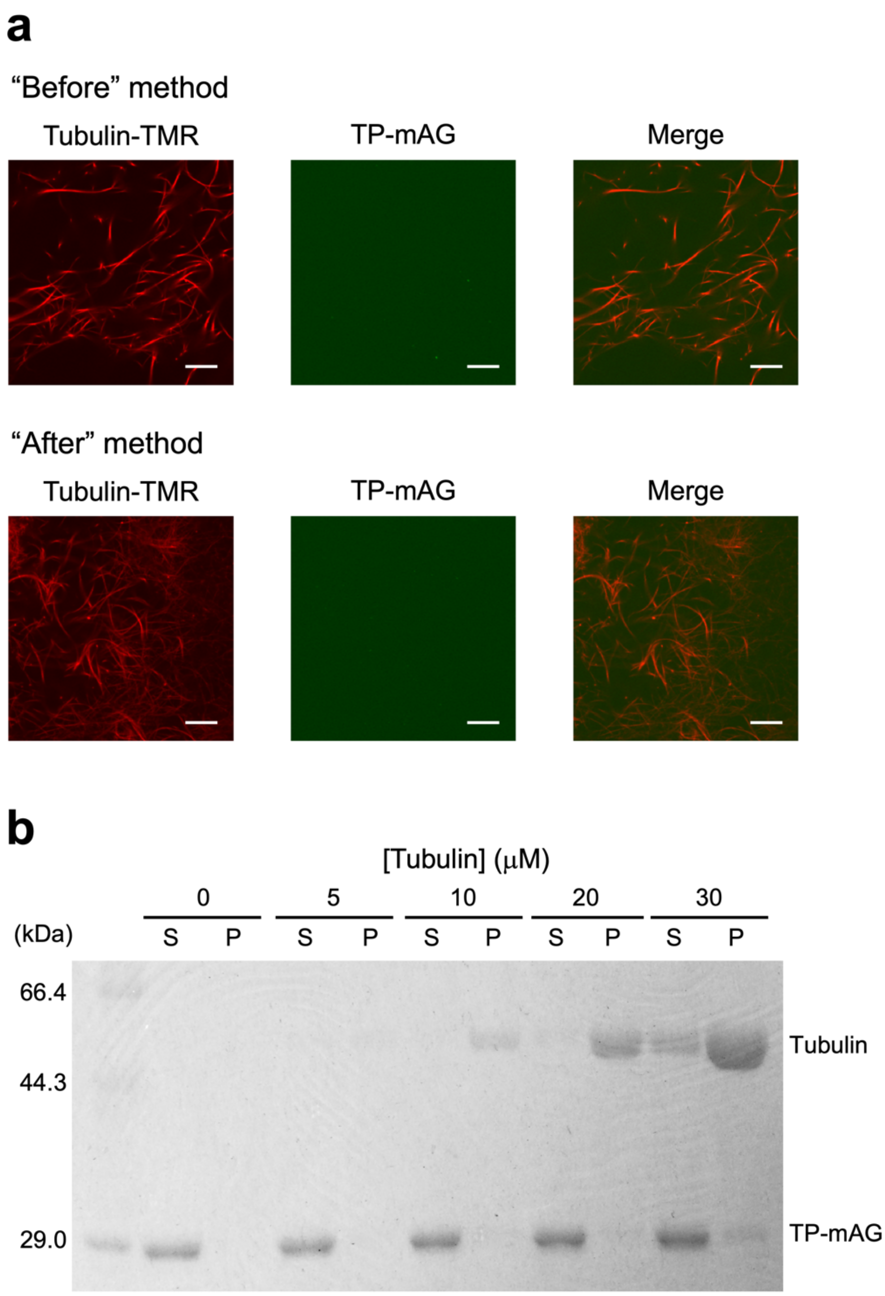


**Supplementary Figure 4.** Binding analysis of TP–mAG to GMPCPP MTs. **(a)** CLSM images of MTs incubated with TP–mAG prepared by the "Before" and "After" methods. Preparation concentrations: [tubulin] = 3.2 μM; [tubulin–TMR] = 0.8 μM; [TP–mAG] = 8 μM; [GMPCPP] = 0.2 mM. Scale bars, 10 μm. **(b)** Binding assay of TP–mAG to MTs. MTs were prepared using 37 μM TP–mAG and various concentrations of tubulins by the "Before" method. SDS-PAGE results of the supernatants (S) and the pellets (P) obtained by centrifugation of the mixture solutions. Most of TP–mAG was not bound to MTs observed as pellets even using high concentration of tubulin.

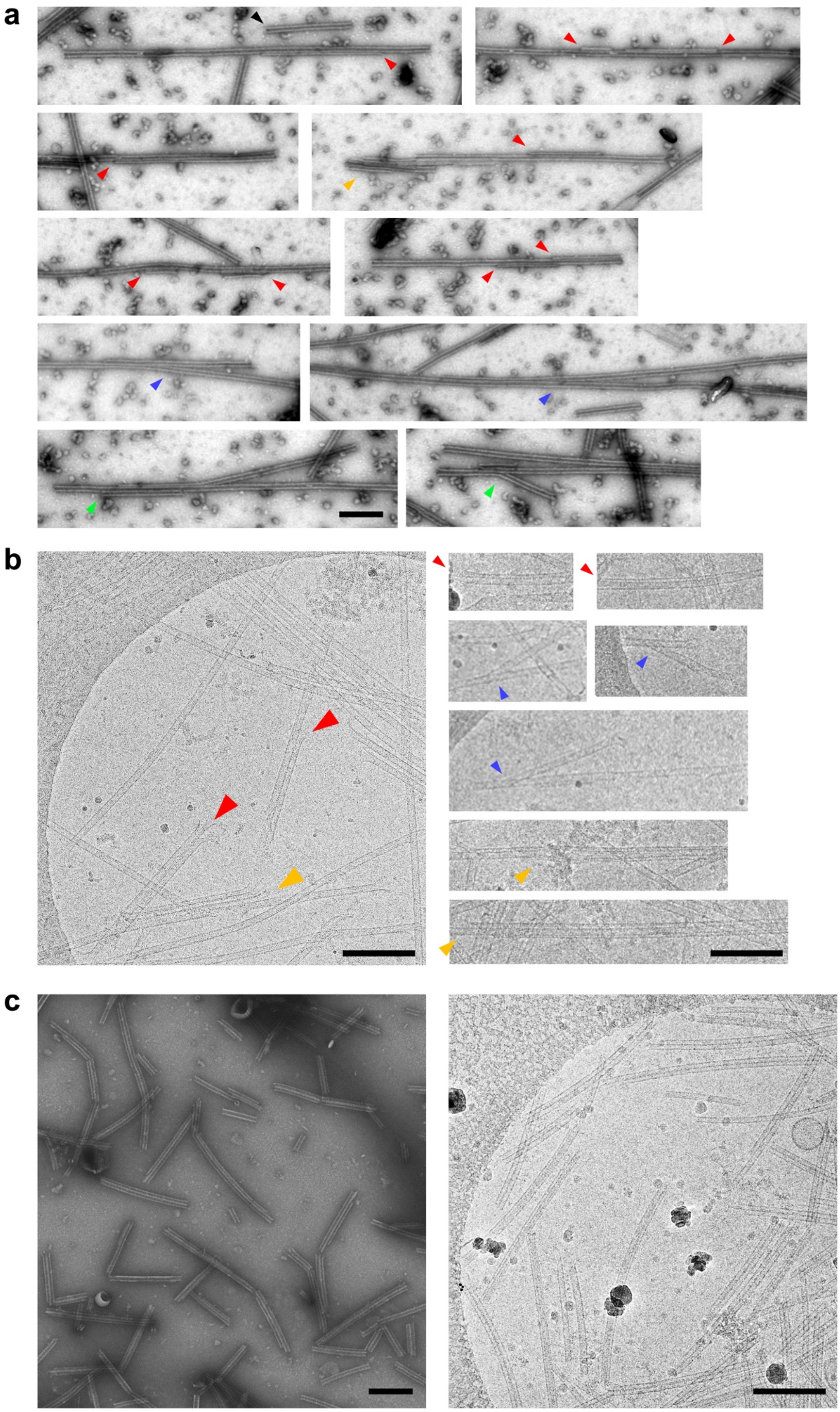


**Supplementary Figure 5.** Data related to formation of doublets, branches, and multiplets. **(a)** Representative negative-stain EM images of TP–AG-incorporated GMPCPP MTs prepared by the "After" method without further incubation. Doublet MTs, branched MTs were essentially same with and without further incubation. Scale bar, 250 nm. **(b)** More representative cryo-EM images of TP–AG-incorporated GMPCPP MTs prepared by the "After" method with further incubation at 37°C for 30 min. Scale bars, 250 nm. Red arrowheads, doublet MTs. Blue arrowheads, branching MTs. Green arrowheads, both doublet and branching. Orange arrowheads, multiplet MTs. **(c)** Negative-stain EM (left panel) and cryo-EM (right panel) images of native doublet MTs purified from Tetrahymena cilia. Tetrahymena doublet MTs were prepared as described previously[1]. In the EM images, there are also some singlet MTs observed due to the breakage of B-MTs because of the sonication of the preparation process. Scale bars, 250 nm.

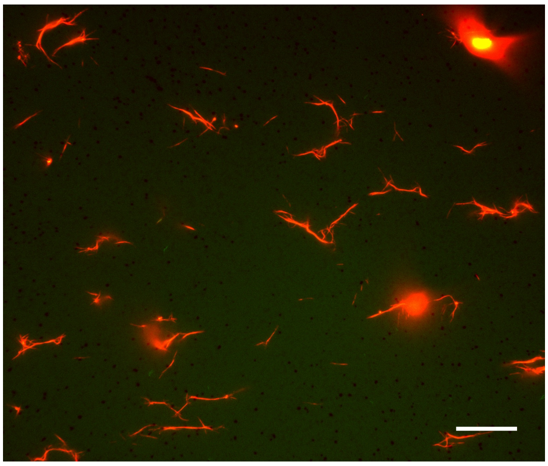

**Supplementary Figure 6.** Fluorescence microscopy image of TP–AG-incorporated GTP MTs prepared by the "Before" method on the kinesin-coated substrate. Preparation concentrations: [tubulin] = 4 μM; [tubulin–TMR] = 1 μM; [TP–AG] = 10 μM; [GTP] = 1 mM. Scale bar, 20 μm.

**Supplementary Movie 1.** A movie that illustrates motility of GMPCPP MTs prepared with TP–AG or buffer by the "Before" method (Fig. 5a). The movie is 100 times faster than the original speed. Scale bar, 20 μm.

**Supplementary Movie 2.** A movie that illustrates motility of MT aster structures formed by TP–AG (Fig. 5c). The movie is 200 times faster than the original speed. Scale bar, 20 μm.
